# Neutralization of variant under investigation B.1.617 with sera of BBV152 vaccinees

**DOI:** 10.1101/2021.04.23.441101

**Authors:** Pragya D. Yadav, Gajanan N. Sapkal, Priya Abraham, Raches Ella, Gururaj Deshpande, Deepak Y. Patil, Dimpal A Nyayanit, Nivedita Gupta, Rima R. Sahay, Anita M Shete, Samiran Panda, Balram Bhargava, V. Krishna Mohan

**Affiliations:** Indian Council of Medical Research-National Institute of Virology, Pune, Maharashtra, India, Pin-411021; Bharat Biotech International Limited, Genome Valley, Hyderabad, Telangana, India Pin-500 078; Indian Council of Medical Research, V. Ramalingaswami Bhawan, P.O. Box No. 4911, Ansari Nagar, New Delhi, India Pin-110029

**Keywords:** VUI, B.1.617, neutralization, BBV152, SARS-CoV-2

## Abstract

The drastic rise in the number of cases in Maharashtra, India has created a matter of concern for public health experts. Twelve isolates of VUI lineage B.1.617 were propagated in VeroCCL81 cells and characterized. Convalescent sera of the COVID-19 cases and recipients of BBV152 (Covaxin) were able to neutralize VUI B.1.617.

## Text

Several variants of the Severe Acute Respiratory Syndrome Coronavirus-2 (SARS-CoV-2) [B.1.1.7(United Kingdom), B.1.351(South Africa), B.1.1.28 (Brazil P1, P2)] have been reported in India during January to April 2021 [1,2]. The daily rise in the reported number of SARS-CoV-2 infections has also reached striking in the country. Among all the affected states, the western State of Maharashtra been hit the hardest during this second wave[3]. The drastic rise in the number of cases in Maharashtra has been an enigma and matter of concern to public health experts. Here, we report the isolation of SARS-CoV-2 of new lineage B.1.617 with several spike mutations from Maharashtra state, India. Further, we investigated the neutralization efficiency of convalescent sera (n=17) and the sera collected from BBV152 (Covaxin) vaccinated individuals (n=28) against the B1(D614G) and B.1.617 variants.

From January to March 2021, we had identified 146 COVID-19 cases from the state of Maharashtra. Nasopharyngeal/oropharyngeal swabs of all the cases were screened with next-generation sequencing using the Illumina platform [4]. Among these 15 retrieved SARS-CoV-2 sequences demonstrated the presence of a combination of L452R and E484Q mutations. Virus isolation was attempted from all the 15 specimens using Vero-CCL-81 cells [5]. Twelve clinical specimens displayed typical rounding of the cells, syncytia formation, and detachment of the cells on the 4th post-infection day (PID). All the twelve virus isolates were passaged, tittered, sequenced, and used further for performing PRNT50 assay as described earlier[6].

A total of 23 non-synonymous changes were commonly observed amongst the retrieved sequence. Out of which, seven conserved non-synonymous changes were observed at spike protein (G142D, E154K, L452R, E484Q, D614G, P681R, Q1071H) concerning the Wuhun-Hu1 sequence (Figure 1 A). Noticeably, the combination of L452R and E484Q mutations were present in many sequences submitted from different countries to the Global Initiative for Sharing Avian Influenza Data (GISAID). Along with these, we marked other conserved mutations as depicted in Figure 1 A. These sequences were classified as B.1.617 using PANGOLIN classification. A phylogenetic analysis was performed on the retrieved sequences along with the representative sequences that demonstrated a separate cluster of B.1.617 lineage. Moreover, three distinct sub-clusters of B.1.617 lineages were observed with mutations in spike first T95I; second H1101D and third V382L, V1175Y (Figure 1 B). One of these sequences (MCL-21-H-741) has an E484K mutation that leads to its separate distinct clustering.

**Figure 1:**
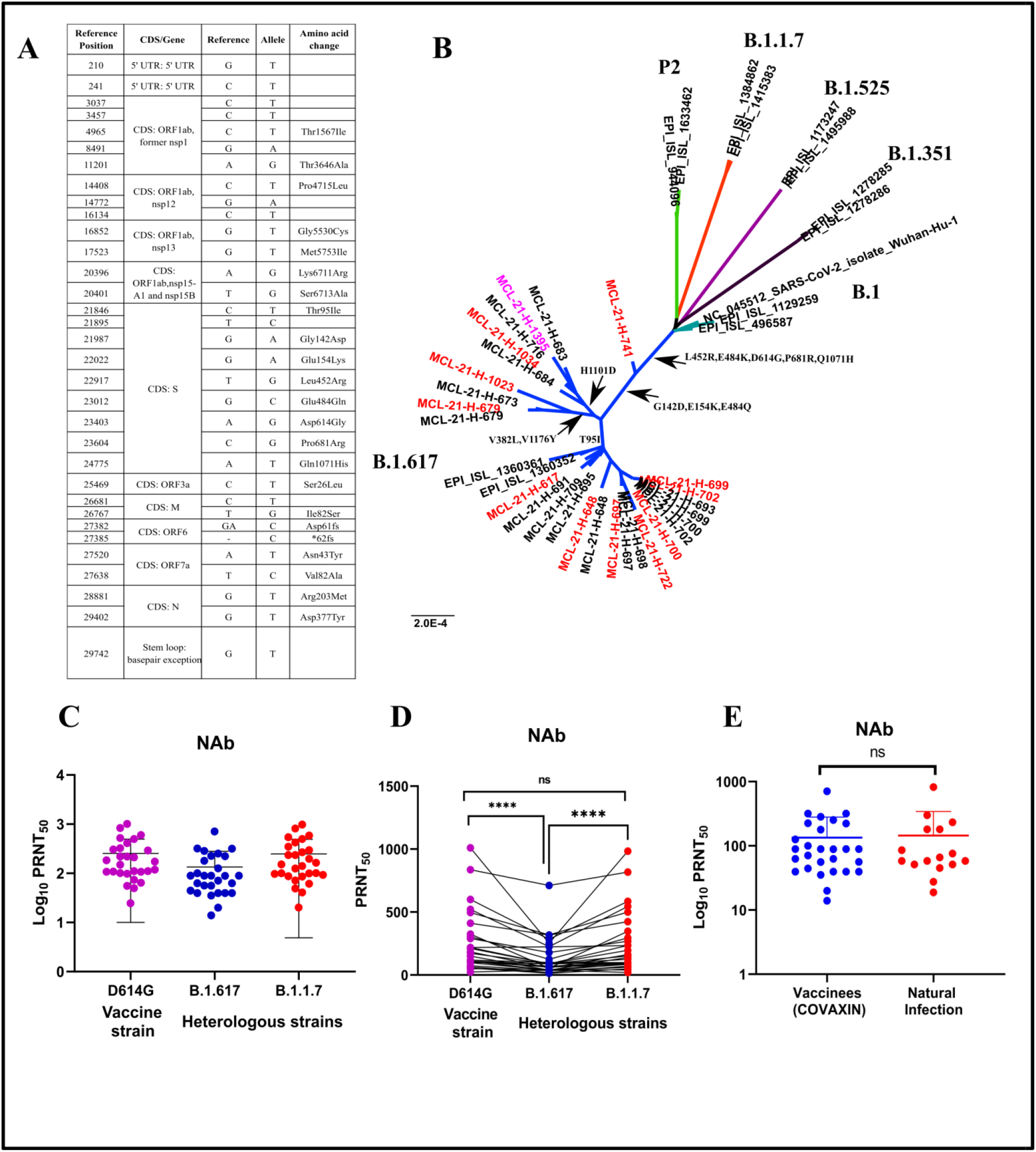
nCharacteristics and neutralization of VUI B.1.617 variant: **A)n**The common nucleotide changes observed in majority of the isolates and clinical sequences. **B)n**A neighbor joining tree was generated using a Tamura 3-paramter model with gamma distribution and a bootstrap relplication of 1000 cycles. Isolates are marked in red colour and sequence from foreign traveller marked in pink colour. The representative sequces from other caldes are represented as B.1.1.7 (red), B.1.617 (blue), B.1.351 (black), B.1.525 (purple) Brazil P2 (light green), and B1 (light blue). Individual spike mutations specific to the clusters are marked using the arrows. **C**) Scatter plot depiting the neutralizing response of the individual sera (n=28) vaccinated with BBV152 (Covaxin) collected during phase II clinical trial for the prototype B1 (D614G) (pink), B.1.1.7 (red), B.1.617 (blue). Red soild line indicates the geometric mean titre and error bar depicts 95% confidence interval. **D)** Neutraliztion of the matched-pair samples compared to prototype D614G (pink), B.1.1.7 (red), and B.1.617(blue). Neutralization reduction by a factor of 1.95 and 1.8 was observed against the B.1.617 variant for B1 (D614G) and B.1.1.7 variant respectively. A reduction factor of 1.06 was observed between the B1 (D614G) and B.1.1.7 variant. A two-tailed pair-wise comparison was performed using the Wilcoxon matched-pairs signed-rank test with a p-value of 0.05. **** represent p value <0.001 and **p value=0.0038, ns= nonsignificant p value. **E)**Neutralization of the COVID-19 recovered cases sera (n=17) of B.1.1.7 (n=2), B.1.351(n=2), B.1.1.28.2(n=2) and B1 lineage (n=11) infected individuals PRNT50 values against B.1.167 variant were compared with vaccine recipient serum samples. A two-tailed pair-wise comparison was performed using the Mann-Whitney test with a p-value of 0.05.

The variant under investigation (VUI) B.1.617 isolates were obtained, from clinical specimens obtained from asymptomatic individuals (ages: 14, 16, 17, 36, 42, 50, and 55 years) and symptomatic cases (ages: 26, 28, 54, 60 and 77 years). The symptomatic cases had mild upper respiratory tract infection including low-grade fever, cough, and sore throat. None of the cases developed the severe disease during infection and recovered completely in due course. One of the virus isolates was from an international traveler who had returned from the United Arab Emirates to India in March 2021.

The neutralization efficacy of the VUI B.1.617 variant was compared with prototype strain B1 (D614G) and B.1.1.7 variant using sera of 28 BBV152 (Covaxin) vaccinated individuals, collected during the phase II clinical trial [7]. For D614G vs. B.1.617, the GMT ratio was 1.95, (95% CI:1.60 - 2.38 and p-value <0.0001) resulting in a statistically difference. Similarly, the GMT ratio comparison of B.1.1.7 was significantly higher than the GMT for B.1.617 (GMT ratio 1.84, 95% CI: 1.50 - 2.27, p value< 0.0001) and the CI was not within the equivalence interval (Figure 1 C and 1D). The comparison of D614G and B.1.1.7 showed equivalent responses with a GMT ratio of 1.06 which is close to 1, and the 95% CI (1.02 to 1.10) was well within the statistical equivalence.

Sera samples collected from COVID-19 recovered individuals (n=17) infected with lineage B.1.1.7 (n=2), B.1.351(n=2), B.1.1.28.2(n=2), and B1 (n=11) were used to perform PRNT50 against B.1. 617 variant, and the results were compared with the results obtained by using sera samples collected from vaccine recipients. The GMT values for vaccine recipients were 88.48 (95% CI: 62.02 to 126.2) and for recovered cases 86.85 (95% CI 52.04 to 144.9). This demonstrated that neutralizing capacity against B.1.617 is similar from sera of vaccinated individuals and that of recovered cases (Figure 1 E).

As any mutations within the amino acid position 438-506 could significantly alter the virus properties leading to enhance infectivity, transmissibility, or escape immunity, both E484Q and L452R amino acid substitutions raised concerns as both were found in the receptor-binding domain (RBD) of the spike protein. This might enhance the transmissibility of the virus and confers resistance to the immune response of the body. It is well known how these mutations behave individually; however, the combined effect of these mutations is still unknown. The studies have suggested that L452R mutation could improve the interaction between spike protein and human angiotensin-converting enzyme 2 (ACE2) receptor [8, 9]. A recent study from California suggested that the newly identified variant of concern (20C/S:452R and 20C/S:452R) with the L452R mutation led to the increased infectivity [8]. Studies have also confirmed that the mutation E484Q could mildly increase the binding affinity to the ACE2 receptor [10]. Both these mutations confer increased binding capacity to ACE receptors which might help the virus to escape the immune response. Further studies are needed to understand the transmissibility and infectivity of the strain. In this study, a drop in neutralization was detected with the B.1.617 variant., However, the reduction of neutralizing capability was limited to 2-fold [11]. Similar results have been observed with sera of BNT162b2 vaccinees which effectively neutralized B.1.1.7 and P.1 variants equally; however, neutralization of B.1.351 variant was reduced but found to be robust [12]. Furthermore, these findings support earlier results comparing BBV152 induced immune responses to neutralize the D614G and B.1.1.7 variants equally [13].

So far 21 countries have detected the B.1.617 variant, most of which have been reported from India [11]. The study’s findings suggest the need for continuing genomic surveillance of B.1.617 and other new emerging variants of SARS-CoV-2. Assessing the clinical efficacy of BBV152 against such variants is underway. It is also important to link genomic surveillance to appropriate public health interventions for preventing massive transmission of the virus. Further studies are needed to understand the transmissibility and infectivity of the B.1.617 variant. Most importantly the assurance of neutralization of B.1.617 variant with sera of BBV152 vaccinees and recovered COVID-19 cases sera will provide the much-needed boost to the COVID-19 vaccination program in India.

## Ethical approval

The study was approved by the Institutional Biosafety Committee and Institutional Human Ethics Committee of ICMR-NIV, Pune, India. under project ‘Molecular epidemiological analysis of SARS-COV-2 circulating in different regions of India’ (20-3-18N).

## Author Contributions

PDY, GNS and PA contributed to study design, data collection, data analysis, interpretation and writing and critical review. RE, RRS, AMS, DYP DAN and GRD contributed to data collection, interpretation, writing and critical review. NG, SP, VKM and BB contributed to critical reviewand finalization of the paper.

## Acknowledgement

Authors gratefully acknowledge the staff of ICMR-NIV, Pune including Mr. Prasad Sarkale, Mr.ShrikantBaradkar, Mr. Rajen Lakra, Mr. VishwajeetDhanure, Dr. Abhinendra Kumar, Mrs. Triparna Majumdar, Mrs. Savita Patil, Ms. Pranita Gawande, Mrs. Kaumudi Kalele, Mrs. Ashwini Waghmare, Ms. Jyoti Yemul, Ms. Manisha Dudhmal, Ms. Aasha Salunkhe and Mr. Chetan Patil for extending excellent technical support.

